# Potential role of *Aphidius ervi* in protecting greenhouse bell pepper cultures from *Myzus persicae* aphids in India

**DOI:** 10.1101/2021.05.03.442443

**Authors:** Ambika Farhat, Ramila Dharmadhrt

## Abstract

Many aphids such as *Myzus persicae* are major agricultural pest against which sustainable solutions must be developed. In the present study, we evaluated the role of *Aphidius ervi (*Hymenoptera: Braconidae), an aphid parasitoid, to control *M. persicae* in bell pepper cultures in greenhouses. Parasitoids were released at one occasion in a greenhouse and M. persicae infested plants were followed before and after parasitoid introduction. We evaluated aphid numbers and parasitism rates at two different controlled temperatures (22°C and 25°C), and we measured diapause levels in released parasitoids. *Aphidius ervi* showed parasitism rates of 10.9±1.2% and 9.1±1.6%, at 21 and 25°C, respectively. The strain of parasitoids we used was not entering diapause under any temperature condition. Therefore, we suggest that *A. ervi* can be further investigated as a protection in greenhouse environments in India. More releases may be necessary to achieve a good protection.

## Introduction

Bell pepper (*Capsicum annuum* L.) is an important vegetable crop grown in India and other parts of the world, where sub-temperate climates are available. Among the many factors identified in limiting successful cultivation of bell peppers, the role of insect pests are extensively reported on this crop (Sanchez et al., 2011). *Myzus persicae* (Sulzer) (Aphididae: Hemiptera) is the most important of them, causing great damage to many crops worldwide. *Myzus persicae* is, for example, causing heavy economic losses in bell pepper in several states in India (Kaur et al., 2004). The problem in greenhouses is that such protected crops provide favorable microclimates for development of insect-pest populations, which limit successful crop production (Sood, 2010).

The aphid can also transmit several viral and fungal diseases in different hosts (Verma et al., 2018). *Myzus persicae* is, for some populations, extremely insecticide resistance (Verma et al., 2018), and consequently extremely difficult to control with pesticides. Therefore, biological control is becoming sustainable approach to manage this pest. Population growth of this aphid is restrained by many natural enemies under field conditions, particularly hymenopteran parasitoids (Starý, 1970; Verma et al., 2018). Augmentative release of hymenopteran parasitoids in greenhouses has been used in different parts of the world (Boivin et al., 2012). On pepper bell, *Aphelinus asychis* Walker, *Aphidius matricariae* Haliday, and *Aphidius ervi* (Haliday) have been identified to parasitize *M. persicae* (Gavkare et al., 2014).

However, little information is available on how these hymenopteran parasitoids respond to different temperatures in terms of parasitism efficiency, and if they express diapause in greenhouse conditions, which would be detrimental to good aphid control. *Aphidius ervi* enters diapause in different parts of the world, such as in Canada or in Europe, but temperatures are colder than in India (Christiansen-Weniger and Hardie, 1999; Polgár et al., 1991; Tougeron et al., 2018a). Assessing diapause and temperature effects in parasitoids is important because they are very likely to be affected by high temperatures associated with climate-change, especially when conditions are already warm such as in sub-temperate areas of pepper bell growth, and in greenhouses (Ghosh and Ballal, 2017; Masry and El-Wakeil, 2020; Tougeron et al., 2019).

## Materials and methods

We surveyed 40 randomly selected bell pepper plants infested with *M. persicae* colonies in two private-own greenhouses (70m^2^ each). One greenhouse was kept at 21°C and the other was kept at 25°C. No insecticide application was done in these greenhouses.

One observation was done on 8 weeks-old plants to count aphids, then parasitoids were released in both greenhouses from our laboratory culture, and one observation was done two weeks after release. Parasitoids were released by groups of 50 (males and females) on 10 random points in each greenhouse, close to the plants. Aphids were visually counted on each plant. Once mummies were formed, they were collected for each plant and kept in petri dishes. Parasitism rate was assessed as the number of mummies / the total number of aphids on the plant two weeks after parasitoid introduction. Mummies were kept until a new parasitoid emerged and diapause rates were evaluated following the protocol of Tougeron et al., (2020); we dissected non-emerged mummies and we classified the content either as a dead parasitoid or a diapausing parasitoid.

## Results

After parasitoid introduction, infestation by *Myzus persicae* decreased by 5.8% in the greenhouse at 21°C, and by 14.7% in the greenhouse at 25°C (Figure 1). Our results indicate that the control at 25°C could be better than at 21°C. However, error bars are overlapping by a lot suggesting that the differences are not major.

**Figure 1:**
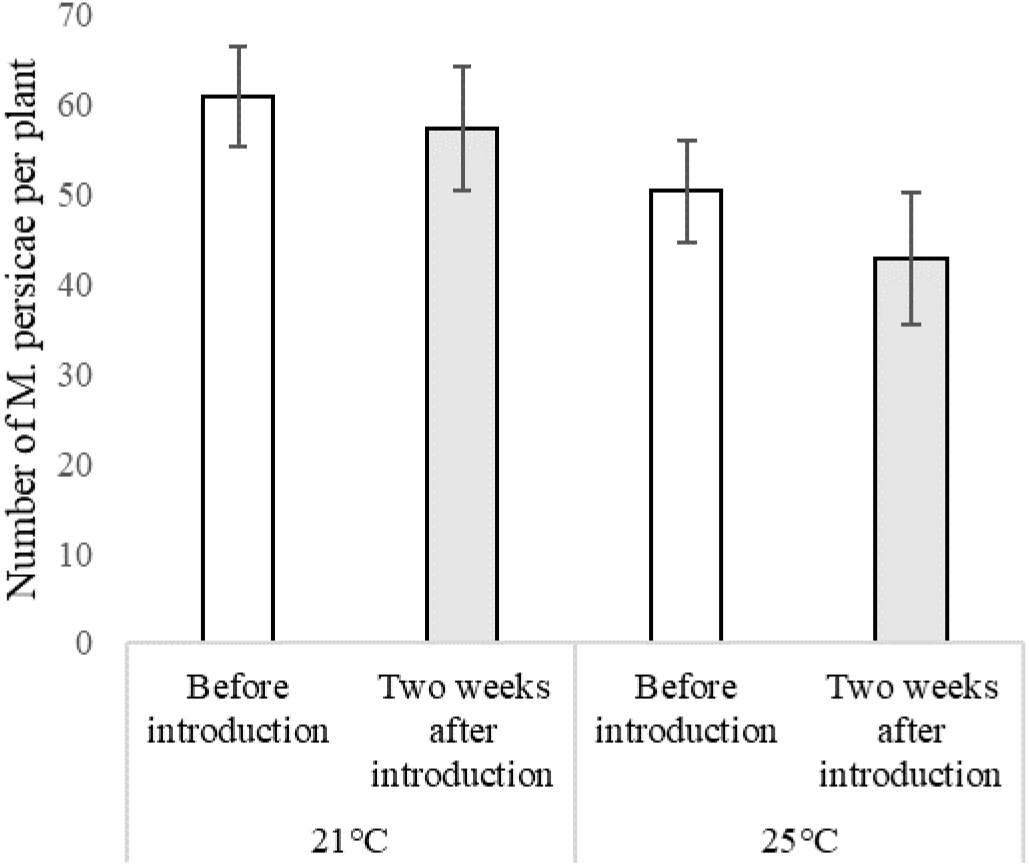
Decrease in mean abundance and standard error of *Myzus persicae* on 20 random plants per greenhouse (at 21°C or at 25°C) after *Aphidius ervi* parasitoid parasitoid release.

Parasitism rates were evaluated at 10.9±1.2% and 9.1±1.6%, at 21 and 25°C, respectively, in greenhouses two weeks after parasitoid release.

Finally, all parasitoids either emerged from their mummies or were found dead when performing the dissections, which indicates that no diapause was found in these released populations, under the temperature conditions we tested.

## Discussion

The present results are promising in terms of control of *M. persicae* under greenhouse conditions. In conjunction with earlier studies (Gavkare et al., 2014; Verma et al., 2018), finding that *A. ervi* can successfully parasitize *M. persicae* in greenhouses is an important piece of information to develop sustainable biocontrol strategies. Some authors that have worked on *A. ervi* also emitted the idea of manipulating sleep in insects in a biological control perspective (Tougeron and Abram, 2017), and we suggest it could be an interesting hypothesis to explore to better control aphids in greenhouses

We did not find major differences between the two temperature treatments. It is know that aphid parasitoids are sensitive to temperature fluctuations in terms of behaviour and physiology (Ismail et al., 2010; Sigsgaard, 2000), but probably temperatures were high enough in our experiments to provide excellent survival to released parasitoids. The parasitoid strain we used was from a population captured earlier in the fields and kept in laboratory environments. It should be noted, however, that many studies have found differences in *Aphidius ervi* parasitism efficiency and life-history traits depending on their origin (Tougeron et al., 2018b, very recently: Saeed et al., 2020).

Further studies are required to develop methods for mass rearing these parasitoids, to better study their behavior against different pest aphids, as well as to develop suitable release techniques for use in Indian greenhouses (Gavkare et al., 2014; Kumar et al., 2020, 2019).

## Notes

### Competing Interest Statement

The authors have declared no competing interest.

